# Constrained Diffusion as a Paradigm for Evolution

**DOI:** 10.64898/2026.03.10.710948

**Authors:** Daniel Lazarev, Anna Sappington, Grant Chau, Ruochi Zhang, Bonnie Berger

## Abstract

A foundational question in computational biology is how to utilize data to describe the forces driving evolution. Here, we view evolution as a novel diffusion process constrained by many biological, physical, and environmental factors affecting organism viability at any given time. We introduce `DiffEvol`, a framework that models evolution as constrained diffusion over a discrete genotype space. Using real-world genomic sequence data alone, `DiffEvol` estimates complex evolutionary constraints by inverting the diffusion dynamics to recover a constrained subspace representing the viable genotype manifold, as well as its evolution over time. Applied to SARS-CoV-2 sequence data from 2020–2024, `DiffEvol` reconstructs constraint functions that recapitulate known viral fitness trends, including a pronounced “phase transition” that occurred following the widespread adoption of the SARS-CoV-2 vaccine. Our constraint subspace representation of the data characterizes such features and trends more clearly. This framework could be used not only to improve forecasting of emergent pathogenic strains, but also to produce more accurate reverse time analyses of their evolutionary dynamics to help identify ancestral variants and the forces having shaped a pathogen’s evolutionary trajectory. More generally, this formulation provides a method for linking observed sequence mutations to an evolving fitness landscape. Thus, constrained subspace diffusion offers a mathematical language for evolutionary dynamics in any system where random variation interacts with slowly-changing structural or global constraints, and can be applied to more complex evolutionary phenomena such as vaccine resistance, viral escape, and protein evolution.

## 1 Introduction

Understanding the factors which contribute to genetic variation over time remains a central challenge in biology. At the molecular level, evolution arises from the interplay between random mutations and selective pressures that determine which variants persist in a population. The recent growth in large-scale genomic sequencing data, particularly during the COVID-19 pandemic, has led to interest in developing quantitative models that can infer evolutionary dynamics directly from observed sequence data.

Recent computational approaches have increasingly relied on statistical and machine learning models to learn evolutionary patterns from large sequence databases [19, 8]. Protein and viral language models have demonstrated impressive predictive power for mutational effects and evolutionary trajectories [13, 14]. However, these black-box predictors generally do not provide interpretability of the mechanisms or constraints that shape evolutionary dynamics. This fundamentally limits their ability to identify more complex forces driving evolution and generalize beyond the observed data distribution.

Here, we take a complementary approach using the simple mathematical language of diffusion, or heat flow, to describe evolution. Specifically, we consider the space of fixed-length genomic sequences and view evolution as driven by random mutations within that space. A key distinction from classical physical diffusion, however, is that most points in sequence space correspond to nonviable or strongly disfavored genotypes. The set of viable sequences can also change over time due to epidemiological, immunological, environmental, or other pressures. Thus, evolution is not an unconstrained diffusion but rather one restricted to a dynamically evolving sequence subspace.

The constraints shaping sequence space can broadly be categorized as either hard constraints, which exclude large fixed regions of nonviable sequence space, and soft time-dependent constraints arising from immune pressure, environmental conditions, and other factors affecting fitness. As the constraints change with time, the accessible evolutionary landscape itself changes, producing trajectories that may never reach steady state on the constrained subspace.

Based on this constrained subspace diffusion modeling approach, we introduce `DiffEvol`, a method that uses observed genomic sequence frequencies, *f*_*t*_, over time to estimate the underlying evolving constraint function, *k*(*t*), which encodes the feasible geometry of the genotype subspace and its evolution. The resulting function offers a unified, interpretable description of evolutionary dynamics that deconvolves the diffusion aspect of evolution, driven by random mutation, from external dynamic constraints, which encode fitness, functional, and epidemiological structure. Using four years of SARS-CoV-2 sequence data from GISAID, we demonstrate how `DiffEvol` recovers evolving fitness constraints and reveals “phase transitions” in viral adaptation, with the potential for both improved predictive forecasting of emergent strains—for example, by leveraging the constraint information into predictive frameworks —and reverse-time reconstruction of ancestral variants.

Although we demonstrate `DiffEvol` using SARS-CoV-2 sequence data as a real-world case study, our framework is not pathogen-specific (see Appendix for results of an application of `DiffEvol` to fifteen years of Influenza A virus data). SARS-CoV-2 exhibits substantial adaptive restructuring under immunological and environmental pressures [8, 2, 24], making it well-suited for illustrating recovered constraint dynamics. However, `DiffEvol` can be applied to any evolving system, such as other viruses as well as to protein evolution in general or other adaptive systems, both biological and non-biological. By modeling these dynamics as diffusion within a time-dependent constraint geometry, `DiffEvol` provides a more interpretable language for studying evolutionary processes and complements existing phylogenetic, statistical, and machine learning approaches to evolutionary modeling.

### Related work

Population genetics has long modeled allele frequency dynamics using stochastic processes. Early theoretical work by Fisher [10, 11] and Wright [25] developed mathematical models of selection, mutation, and genetic drift that describe how allele frequencies change in finite populations. Later work by Kimura [17, 18] showed that in the limit of large population size with weak mutation and selection, these discrete stochastic processes converge to continuous diffusion on allele frequencies. The resulting Wright-Fisher diffusion is still widely used today and remains a central framework in theoretical population genetics.

Our formulation shares important similarities with these classical population genetics approaches. The underlying mathematical structure of `DiffEvol` parallels diffusion-based approaches used in population genetics. However, `DiffEvol` differs in how selective forces are represented and inferred. While traditional formulations model selection via explicit drift terms or fitness coefficients assigned to genotypes, `DiffEvol` represents evolutionary pressures through a time-dependent constraint function estimated as a constraint matrix. Thus, rather than specifying selection coefficients directly, this constraint function acts multiplicatively on the mutation-driven diffusion process, restricting probability mass to a viable genotype subspace. Conceptually, this can be viewed as defining the geometry of the viable genotype manifold at each time point. The constraint-matrix representation provides an interpretable description of how the feasible region of genotype space changes over time, helping to enable both forward prediction and backward reconstruction of evolutionary dynamics.

More recent work has also explored diffusion limits of evolutionary processes. For instance in adaptive dynamics, models have described how discrete mutations converge to a continuous diffusion process in the small-mutation limit [3]. Related ideas appear in data-driven settings: diffusion maps and related spectral methods [4] have shown how geometric structure and low-dimensional manifolds can emerge from local random-walk dynamics, while diffusion-based graph learning approaches [12] provide practical means of capturing global constraints from local transition information. Similarly, recent methods for Bayesian gene constraint estimation [30] infer selective limits with a statistical model using data such as mutational burden and depletion patterns, a conceptual analog of the constraint function estimated here. At a broader physical level, the evolution of such constrained systems parallels gradient-flow formulations of free-energy minimization, as in the Jordan–Kinderlehrer–Otto framework [15, 16], and thermodynamic perspectives on replication and selection [9].

Network representations of genotype spaces have also demonstrated that their connectivity profoundly affects molecular evolution and extinction [1, 27], while network-propagation models have highlighted how local perturbations in mutational landscapes can amplify through gene and protein networks [7]. High-resolution studies of mutation rates and constraint landscapes across the human genome and tumors [22], as well as information-theoretic analyses of large biological datasets [26], underscore the growing role of quantitative constraint modeling in understanding evolution. In the viral context, recent work on SARS-CoV-2 and other pathogens has leveraged such ideas to predict future variants [8], map mutational drivers [13], and forecast viral escape from immune or therapeutic pressures [23].

We emphasize that `DiffEvol` is complementary to existing approaches for modeling evolutionary dynamics. Classical phylogenetic and phylodynamic models describe lineage relationships and population-level parameters over time [19], while fitness-landscape inference methods estimate selection coefficients from mutational patterns [30]. More recently, protein and viral language models have demonstrated impressive predictive power by learning high-dimensional genotype–phenotype mappings directly from sequence data [13, 14, 23]. In contrast, `DiffEvol` focuses on recovering a mechanistically interpretable, time-resolved constraint geometry by inverting mutation-driven diffusion. Its goal is not to replace predictive models, but to provide an invertible dynamical framework that deconvolves mutation from evolving structural constraints and offers a mathematical language for understanding the forces shaping evolutionary trajectories.

## 2 Methods

### 2.1 Background for DiffEvol

For a sequence of nucleotides of length *L*, the *genotype space* is the discrete set

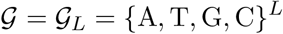

representing all possible length *L* sequences of DNA; we omit the subscript *L* when unambiguous. This alphabet can be modified for RNA or amino acid sequences as needed. We define the constrained genotype subspace, 𝒢_*c*_ ⊂ 𝒢, corresponding to the biologically feasible sequences, given all internal (e.g., biochemical) and external (e.g., epidemiological, immunological, environmental) constraints on the genotype space. These constraints are represented by a *constraint function*

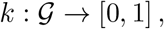

which assigns to each *g* ∈ 𝒢 a value indicating the degree to which it belongs to the feasible subset [28, 21]. Intuitively, *k* encodes the equilibrium or limiting distribution of genotypes for the current environment, while the entropy of *k* quantifies the effective (logarithmic) volume of 𝒢_*c*_ [5, 6]. *We can alternatively view* 𝒢_*c*_ as the support of the constraint function.

We posit that evolution can be viewed as a diffusion process restricted to 𝒢_*c*_. Random mutations drive the diffusive motion across genotype space, but the geometry and topology of 𝒢_*c*_, as determined by the constraint function, govern the accessible directions and the long-term behavior of this diffusion. These geometric features, in turn, emerge from the combined effects of molecular fitness landscapes, interactions with phenotypes and environments, population-level epidemiological pressures, and potential self-interactions among genes.

Because biological and environmental conditions change, the constraint function itself evolves over time. Let *k*(*t*)_*t*∈[0,*T* ]_ denote the family of time-indexed constraint functions, each defining the geometry of the feasible subspace 𝒢_*c*_(*t*) at time *t*. Given empirical frequency data that allow inference of *k*(*t*) for *t* ∈ [0, *T* ], one can estimate the most likely constraint function at a future time. The resulting evolutionary process is therefore a diffusion on a dynamic manifold 𝒢_*c*_(*t*), permitting both forward-time prediction of emergent variants and reverse-time reconstruction of ancestral ones.

This theory assumes (1) a constant mutation rate equal to the mean per-site rate, (2) random and independent mutations, and (3) fixed sequence length *L* across variants. In practice, these assumptions can be relaxed; for example, by introducing a blank or gap character as a fifth nucleotide state or by replacing the Hamming distance with the Levenshtein distance to account for insertions and deletions.

We emphasize that a constant mean per-site mutation rate is not essential to our framework, but is assumed here to simplify applications to real data where per-site mutation rates are rarely available. This assumption does not impose a molecular clock at the level of observable dynamics. All mutation rate behaviors, including acceleration, deceleration, and punctuated evolutionary shifts, are captured by the time-varying constraint function *k*(*t*) inferred from data. In this way, `DiffEvol` explicitly decouples stochastic mutation from time-varying selective constraints. As we explicitly note in the subsequent section, the `DiffEvol` framework naturally generalizes to time- or site-dependent mutation rates.

### 2.2 Constrained Subspace Diffusion

Let *m*_*ij*_ be the mutation rate between *g*_*i*_ and *g*_*j*_, where *g*_*i*_, *g*_*j*_ ∈ 𝒢, and *d*(·, ·) be the Hamming distance. We let **f**_*n*_ ∈ ℝ^*G*^ be the normalized frequency vector at discrete time *n* ∈ [0, *T* ], where *G* = |𝒢|. Although *G* = 4^*L*^ is astronomically large for realistic viral genomes, the set of biologically and epidemiologically viable sequences is orders of magnitude smaller [8], with evolution effectively confined to this much smaller genotype subspace.

#### Discrete-time mutation dynamics

We define the mutation adjacency matrix, *M*, with entries given by

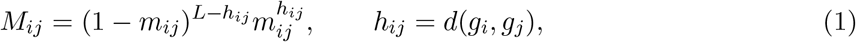

which is symmetric because *h*_*ij*_ = *h*_*ji*_. Assuming a constant per-site mutation rate *m*, we can rewrite Eq. (1) as

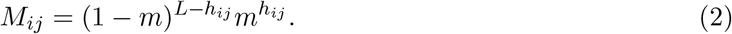

The mathematical derivations below and in the Appendix are equivalent regardless of whether we use Eq. (1) or (2). Next, the degree matrix *D* = diag(*M* **1**_*G*_), where **1**_*G*_ is the column vector of ones of size *G*, yields the row-stochastic transition matrix

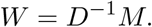

The *unconstrained* discrete diffusion equation, or random walk with transition matrix, *W*, is given by the difference equation

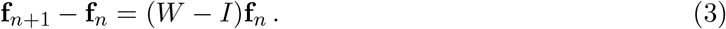

Its solution is given by

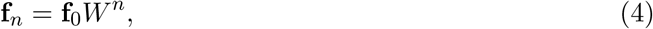

which diffuses mass over genotype space at a rate determined by *m* and converges to the uniform distribution. We note that in the limit of step size approaching zero, Eq. (3) becomes the standard heat equation (see, *e*.*g*., [20])

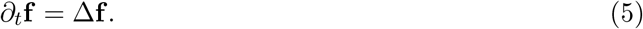

#### Continuous-time embedding and the heat kernel

The discrete-time semigroup 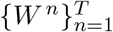 admits a natural continuous-time extension. If we view each replication cycle as a time step Δ*t*, then

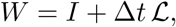

for some generator ℒ. With *t* = *n*Δ*t*, we take Δ*t* → 0, or equivalently, *n* → ∞, which gives

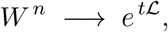

where {*e*^*t*ℒ^}_*t*∈[0,*T* ]_ is the continuous-time mutation-diffusion semigroup. Thus, both *W* ^*n*^ and *e*^*t*ℒ^ represent the same mutation dynamics under either discrete or continuous time, respectively.

The singular value decomposition (SVD) of *W* = *USU* ^⊤^ gives

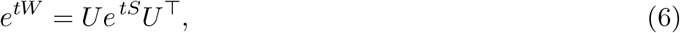

mirroring classical diffusion maps and heat-kernel methods. This provides a continuous-time, smoothed mutation operator that remains stable for large *t* and noisy data.

#### Time-dependent constraints

Evolution does not explore all of 𝒢 but instead remains confined to a dynamically evolving constrained subspace:

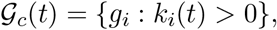

encoded by a constraint function

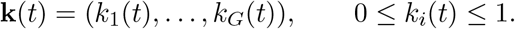

Hard constraints correspond to permanently nonviable sequences (*k*_*i*_(*t*) = 0), while soft constraints vary with biological, epidemiological, or environmental pressures [21, 28, 29]. The constraint function is often considered to evolve on a slower timescale than mutation, which allows mutation and constraint dynamics to be separated.

#### Discrete constrained subspace diffusion

Under constraints, the observed distribution at time *n* is modeled as

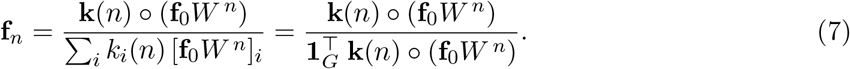

Here **f**_0_*W* ^*n*^ represents diffusion under mutation alone, and the element-wise product with **k**(*n*) restricts this distribution to viable genotypes before renormalization. If **k**(*n*) is constant over time, the sequence converges to a stationary distribution proportional to **k**. If **k**(*n*) evolves, diffusion occurs on a time-varying, slowly drifting constraint manifold 𝒢_*c*_(*n*).

#### Continuous-time constrained subspace diffusion

Using the continuous semigroup *e*^*t*ℒ^, the analogous continuous-time model is

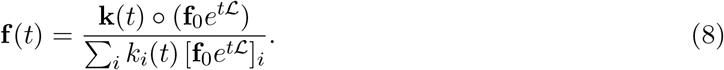

As in discrete time, **k**(*t*) selects the viable region of genotype space, and the heat-kernel operator *e*^*t*ℒ^ propagates mutation continuously. The two formulations are interchangeable under the standard discretization *t* = *n*Δ*t*, with the continuous formulation used for its smoothing effects and favorable properties in the numerical implementation of the method.

Instead of solving the usual heat equation, Eq. (5), Eq. (8) is the solution to the *constrained subspace diffusion equation*,

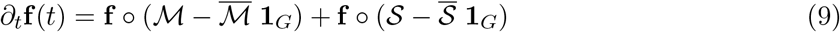

where ℳ = (ℒ **f**_0_*e*^*t*ℒ^) ⊘ (**f**_0_*e*^*t*ℒ^) represents the contribution due to the random mutations, while 𝒮 = *∂*_*t*_**k**(*t*) ⊘ **k**(*t*) represents that due to the evolving constraints, with 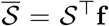 and 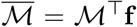 their mean values taken with respect to the frequencies. Eq. (9) reduces to Eq. (5) if **k**(*t*) = **1**_*G*_.

(See Appendix for the derivation of Eq. (9).)

### 2.3 The DiffEvol Method

Having formulated mutation as a diffusion on a constrained submanifold of genotype space, we now describe how this framework yields a practical method for learning the time-evolving constraint functions directly from observed frequency data. The key idea is that the constrained subspace diffusion model can be *inverted* : given the observed frequencies {**f**_*n*_} and the mutation kernel *W*, we can solve for the constraint function **k**(*n*) at each time point. The resulting time series **k**(0), …, **k**(*N*) encodes the geometry of the dynamically evolving feasible subspace 𝒢_*c*_(*n*).

#### Direct inversion in discrete time

The discrete constrained subspace diffusion equation, given by Eq. (7), implies the proportionality **k**(*n*) ∝ **f**_*n*_ ⊘ (**f**_0_*W* ^*n*^), with ⊘ denoting elementwise (Hadamard) division. Normalizing yields,

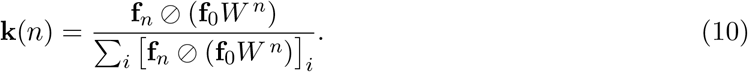

Computing Eq. (10) for each *n* ∈ [0, *N* ] produces the *constraint matrix K* ∈ ℝ^*N ×G*^, whose *n*th row describes the feasible region of genotype space at time *n*.

#### Continuous-time analogue and the heat kernel

Mutation can also be modeled in continuous time via the semigroup *g*(*t*) = **f**_0_*e*^*tW*^, which is the solution to the continuous time and discrete space diffusion heat equation, *∂*_*t*_*g*(*t*) = *Wg*(*t*). The constrained continuous-time equation, given by Eq. (8), implies, exactly as in the discrete case, that **k**(*t*) ∝ **f** (*t*) ⊘ **f**_0_*e*^*tW*^, and thus the continuous-time inverse solution is given by,

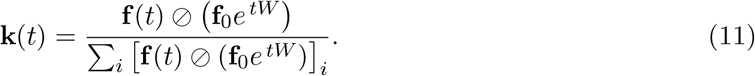

In practice, continuous-time inversion is computed spectrally, using the SVD formulation in Eq. (6). This gives a smoothed, continuous-time mutation operator that is well-conditioned for large *t*.

#### Estimating the temporal dynamics of constraints

The sequence 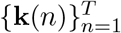 typically evolves on a slower timescale than mutation. We can therefore treat the rows of *K* as a time series that lies near a low-dimensional submanifold in ℝ^*G*^. This makes possible:

- **Forward prediction:** estimating future constraint vectors **k**(*T* +1), **k**(*T* +2), … and thereby future prevalence distributions via Eqs. (7) or (8).
- **Backward reconstruction:** inferring likely ancestral constraint states and ancestral prevalence distributions, provided the explored region of *W* is invertible.
- **Quantifying selective shifts:** identifying abrupt or gradual changes in **k**(*n*) corresponding to selective sweeps, immune or therapeutic pressures, or major epidemiological transitions.

#### Summary

The `DiffEvol` method is a principled inversion of an underlying constrained subspace diffusion process. It decouples mutation from constraint, reconstructs a time-resolved viability landscape from empirical frequencies, and provides a coherent forward–backward dynamical model for predicting future variants and inferring ancestral ones. In this sense, **k**(*n*) is not merely a fitted parameter but a meaningful description of the evolving constraint geometry that steers viral evolution.

## 3 Results

### 3.1 Toy model

To test the performance of our method and its underlying assumptions, we first built a toy model. Let *K*_true_ be the ground truth constraint matrix, *F*_*n*_ the frequency matrix with added noise, and *K*_est_ the estimated constraint matrix using `DiffEvol`. We added noise drawn from a normal distribution 𝒩 (0, 10^−5^), where the standard deviation was chosen so that it is approximately two orders of magnitude smaller than the mean value of the frequency matrix entries.

To better simulate real data, and to test our use of the mean mutation rate in the transition matrix used to estimate the constraint matrix (Methods), we used a variable mutation rate in the ground truth transition matrix used to generate the simulated frequencies. We generated a vector of *G* mutation rates by taking the absolute value of numbers drawn from 𝒩 (*m, m*). We used *m* = 10^−5^ for the simulations shown in plots in Fig. 2 and Table 1.

**Figure 1:**
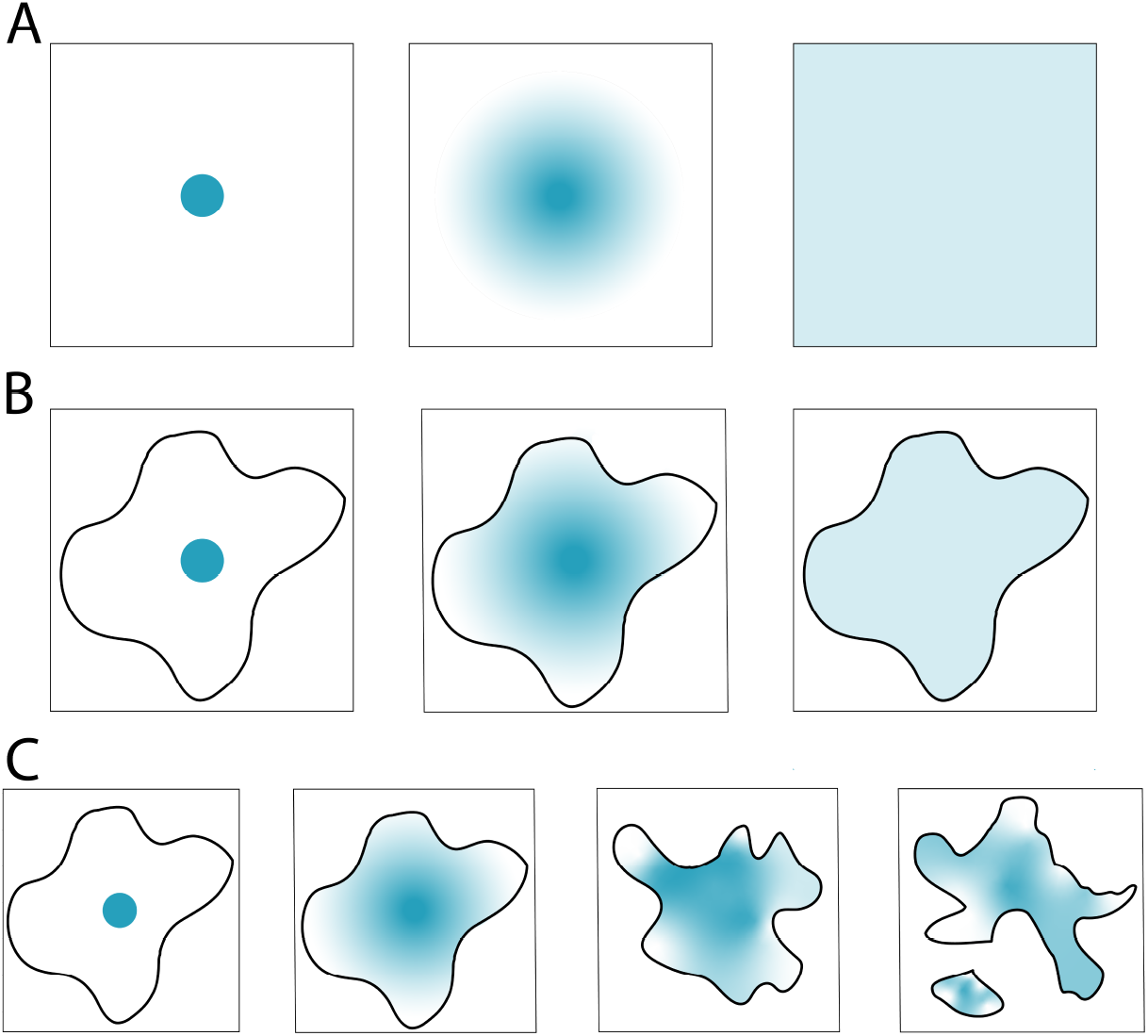
Schematic illustrating the diffusion of a ball of gas for three different constraint conditions. **(A)** Unconstrained diffusion. **(B)** Constrained subspace diffusion. **(C)** Constrained subspace diffusion with evolving constraints.

**Figure 2:**
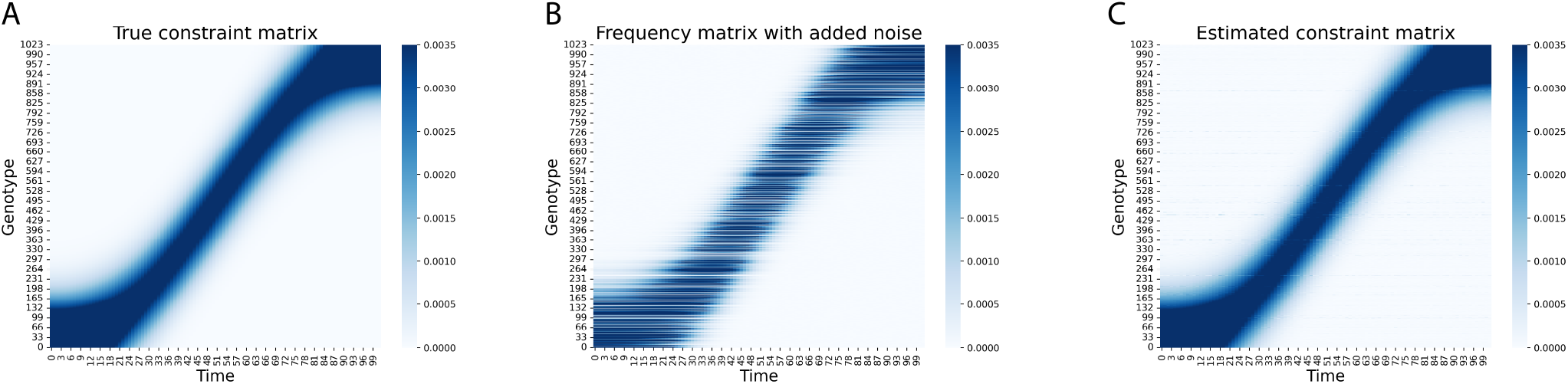
**(A)** The ground truth constraint matrix, *K*_true_. **(B)** The frequency matrix, *F*_*n*_, with added noise. **(C)** The estimated constraint matrix, *K*_est_, using `DiffEvol`.

**Figure 3:**
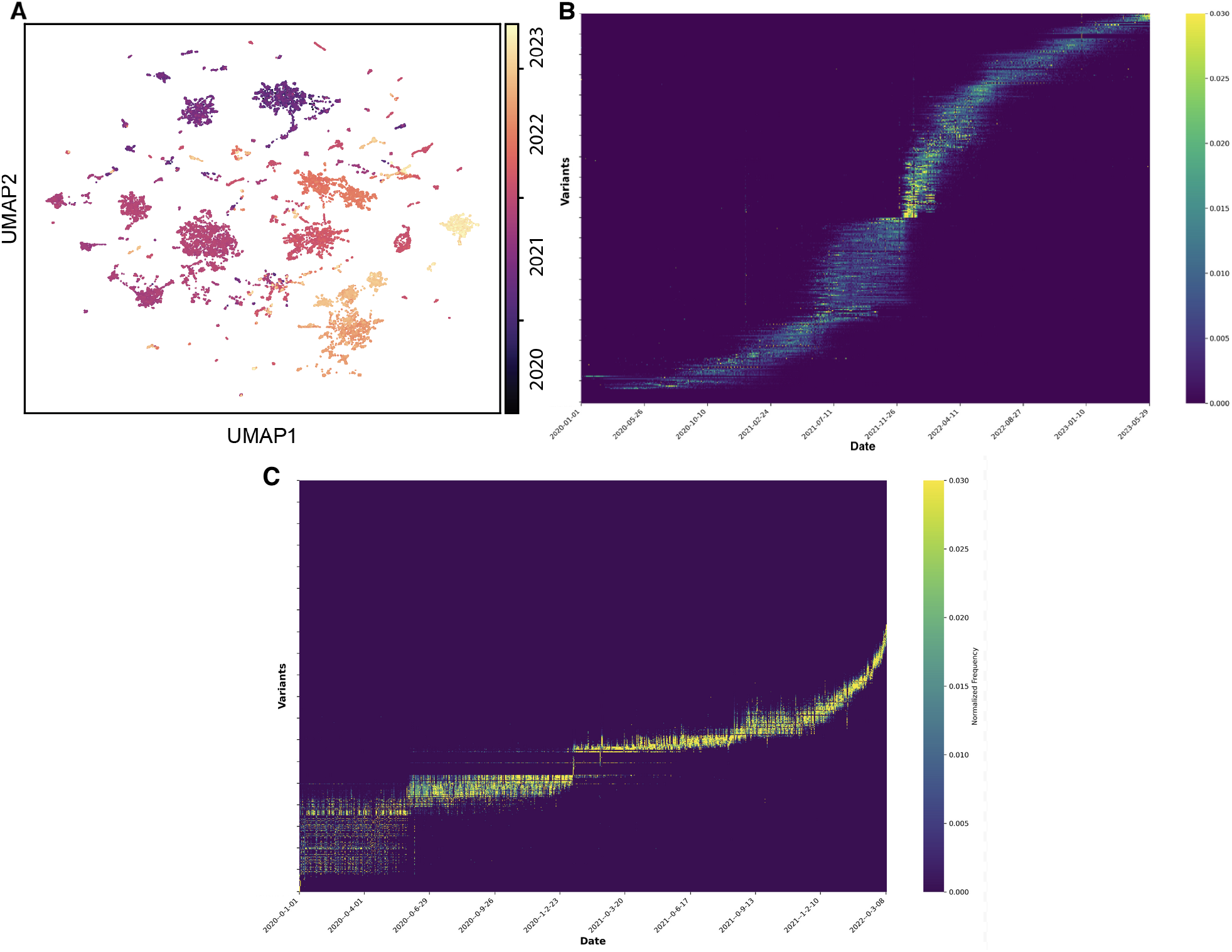
**(A)** UMAP plot and **(B)** heat map based on frequency data of the SARS-CoV-2 virus nucleotide genotypes over three years. **(C)** Corresponding constraint matrix using `DiffEvol`. Variants are ordered by their mean emergence time.

**Figure 4:**
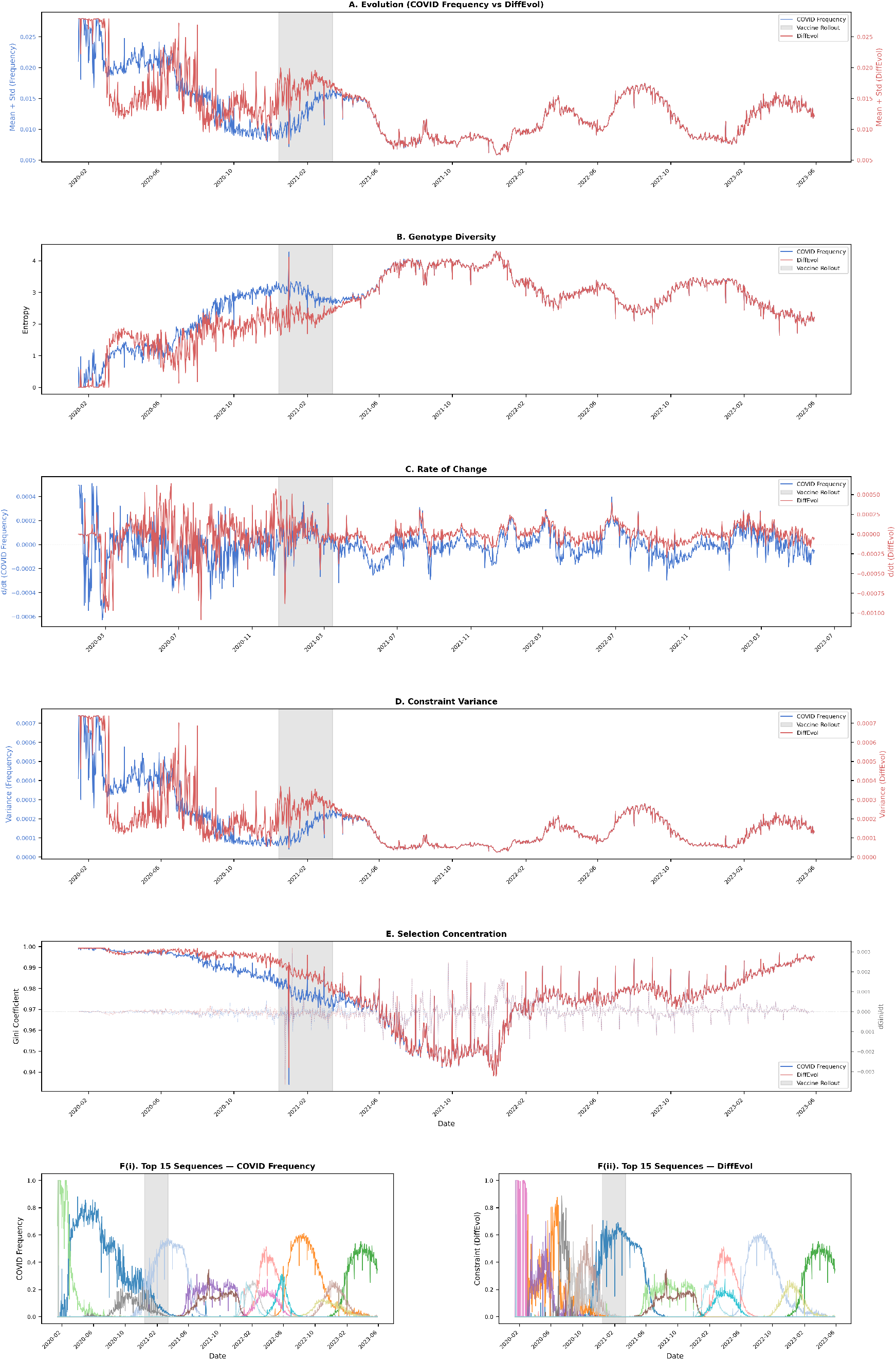
Comparison of raw frequency dynamics and DiffEvol-derived constrained subspace diffusion for SARS-CoV-2 (2020–2023). `DiffEvol` denoises raw frequency data (blue) into interpretable constraint dynamics (red), revealing punctuated increases in evolutionary constraint around the vaccine rollout (gray). **(A–E)** show smoother, biologically meaningful trends in constraint intensity, diversity, rate of change, variance, and selection concentration compared to raw data. **(F i–ii)** illustrate that `DiffEvol` compresses thousands of sequences into a small set of coherent evolutionary trajectories capturing dominant adaptive modes.

**Table 1:**
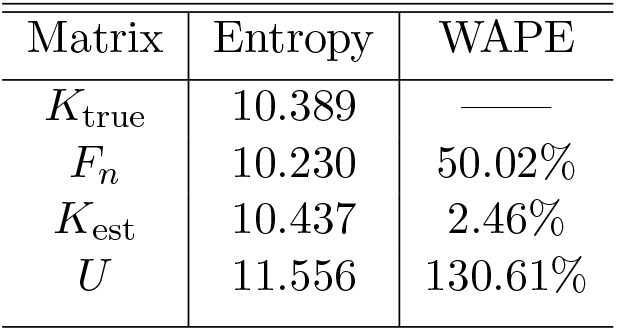
Performance of `DiffEvol` in reconstructing the constraint matrix from the frequency matrix with added noise. WAPE was measured relative to *K*_true_ and *U* is the uniform matrix with entries given by 1/*G* for which the entropy attains its maximum. Used variable mutation rates in generating the frequency matrix.

| Matrix | Entropy | WAPE |
| --- | --- | --- |
| $K_{\text{true}}$ | 10.389 | — |
| $F_n$ | 10.230 | 50.02% |
| $K_{\text{est}}$ | 10.437 | 2.46% |
| $U$ | 11.556 | 130.61% |

Note that the WAPE for $U$ in Table 1 is greater than 100% because of the large number of entries for which $U_{ij} > K_{\text{true}}$ .

To assess the performance of our method on simulated data, we use the weighted absolute percent error (WAPE), given by

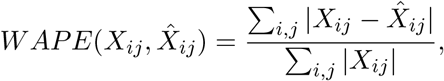

where *X*_*ij*_ is the real data and 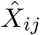 is the predicted data.

Note that the WAPE for *U* in Table 1 is greater than 100% because of the large number of entries for which *U*_*ij*_ > *K*_true_.

### 3.2 SARS-CoV-2 model

#### Data acquisition and preprocessing

SARS-CoV-2 genomic sequences were obtained from the GISAID database from dates spanning January 1, 2020 to May 29, 2023. FASTA files were parsed to extract sequence data, collection dates, and geographic locations. For each record, we extract the collection date, location, and full nucleotide sequence. Unique sequences were identified and assigned sequential indices, and this resulted in a map between the original 47,550 filtered sequences and a compressed set of 3,262 unique variants. This alignment step guarantees that sequences are matched with their corresponding temporal frequency profiles.

#### Occurrence matrix

Temporal occurrence patterns were encoded in a sparse matrix where rows represent time points (823 unique dates) and columns represent sequence variants. Sequences were filtered based on observational coverage, retaining variants with at least 10 total occurrences across the study period. This removed rare or potentially artifactual sequences while preserving biologically relevant variants.

#### Frequency normalization

Raw occurrence counts were converted to normalized frequencies through a two-stage process. First, we normalized counts by time point (column-wise) and compute the relative frequencies that sum to 1 at each time point. Next, frequencies were normalized by variant (row-wise) to standardize the temporal profile of each variant across time. This ensured that variants with different absolute abundances could be compared on a similar scale.

To denoise the raw frequency data and extract underlying evolutionary constraints, we apply `DiffEvol`, which models evolution as constrained subspace diffusion over the discrete genotype space defined by our dataset. The constraint matrix *K* consists of *T* = 823 time points and *G* = 3, 262 genotypes. For comparison, we report the following metrics for both the raw frequency matrix and the constraint matrix *K*:

1. **Mean Evolution**. Temporal trajectories of mean constraint or frequency values (averaged across all genotypes at each time point). This depicts overall population-level dynamics and responses to selective pressures.
2. **Entropy**. Shannon entropy of the distribution at each time point, quantifying genotypic diversity. The effective number of genotypes was computed as the inverse Simpson index: 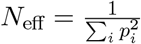, where *p*_*i*_ represents normalized frequencies or constraints.
3. **Diversity**. Number of dominant variants at a given time.
4. **Rate of Change**. First-order temporal derivatives (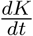 and 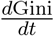) to identify periods of rapid evolutionary change and potential phase transitions.
5. **Gini Coefficient**. Measures concentration of the frequency or constraint distribution, with values near 1 indicating dominance (high concentration) by few variants and values near 0 indicating uniform distribution (random chance). Temporal changes in the Gini coefficient 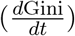 depict shifts in competitive dynamics.
6. **Variance**. Temporal variance across genotypes, indicating the spread of the distribution and the stability of the constraint landscape.
7. **Top** *n*(= 15) **sequence constraints**. Depicts which *n* variants have the highest mean constraint (best “fitness” on average).

To examine the impact of vaccination on viral evolution, we identified the vaccine rollout period (December 15, 2020 to March 15, 2021) and compared pre-vaccine, vaccine rollout, and post-vaccine periods. Statistical distributions of constraints before and after this period were compared to quantify the shifts in the evolutionary landscape. Comparison of 4 shows that `DiffEvol` is effective at extracting evolutionary signals from noisy observational data. Notably, the vaccine rollout period marked a clear evolutionary regime shift and directional selection where previously successful variants were systematically eliminated (F) and the selective sweep in the middle of the vaccination period converged rapidly to low diversity (E) with the dominance of a single variant. This pattern is consistent with strong immune selection following vaccination that overwhelms other evolutionary pressures (transmission fitness, geographic effects, stochastic drift).

Finally, we also applied `DiffEvol` to Influenza A virus (Subtype H1N1). The corresponding plots depicting the frequency and constraint matrices can be found in Appendix.

## 4 Discussion

We introduced `DiffEvol`, a framework that models evolution as a diffusion process constrained by complex, time-varying molecular, immunological, environmental, and epidemiological forces. By inverting the diffusion dynamics on genotype space, `DiffEvol` infers the evolving constraint functions that reflect the various forces driving evolutionary change. Applied to SARS-CoV-2, it reconstructs constraint shifts corresponding to major fitness transitions, including those following vaccine rollout.

In future work, we hope to explicitly analyze both forward-time forecasting of variants and reverse-time reconstruction of ancestral states using `DiffEvol`’s constraint functions. We also aim to enable the incorporation of multiple data modalities and multiscale data (e.g., cell-level, protein, and gene expression data) to further inform the model.

Taken together, our formulation unifies stochastic mutation and structured selection within a single diffusion–constraint framework, providing an interpretable alternative to black-box predictive models. It offers a general mathematical language for adaptive processes where random variation interacts with evolving structural constraints, from viral evolution to molecular and cellular systems.

## Acknowledgments

*Funding acknowledgment*. This work was supported by NIH grants R35GM141861 (to BB) and T32GM14427 (to Harvard/MIT MD-PhD Program) from the National Institute of General Medical Sciences and a Hertz Foundation Fellowship (AS). The content is solely the responsibility of the authors and does not necessarily represent the official views of the National Institute of General Medical Sciences, the National Institutes of Health. The authors declare no known conflict of interest.

## Notes

### Competing Interest Statement

The authors have declared no competing interest.

### Summary of Updates

Introduction and Related Work sections updated.

## References

[1] Aguirre, J., Catalán, P., Cuesta, J., Manrubia, S.: On the networked architecture of genotype spaces and its critical effects on molecular evolution. Open Biol. 8, 180069 (2018)

[2] Bradley, C.C., Wang, C., Gordon, A.J.E., Wen, A.X., Luna, P.N., Cooke, M.B., Kohrn, B.F., Kennedy, S.R., Avadhanula, V., Piedra, P.A., Lichtarge, O., Shaw, C.A., Ronca, S.E., Herman, C.: Targeted accurate rna consensus sequencing (tarc-seq) reveals mechanisms of replication error affecting sars-cov-2 divergence. Nature Microbiology 9(5), 1382–1392 (2024)

[3] Champagnat, N., Lambert, A.: Evolution of discrete populations and the canonical diffusion of adaptive dynamics. The Annals of Applied Probability 17(1), 102 – 155 (2007)

[4] Coifman, R.R., Lafon, S., Lee, A.B., Maggioni, M., Nadler, B., Warner, F., Zucker, S.W.: Geometric diffusions as a tool for harmonic analysis and structure definition of data: Diffusion maps. Proceedings of the National Academy of Sciences 102(21), 7426–7431. 10.1073/pnas.0500334102

[5] Coifman, R.R., Wickerhauser, M.V.: Entropy-based algorithms for best basis selection. IEEE Transactions on Information Theory 38(2), 713–718 (1992)

[6] Cover, T., Thomas, J.: Elements of Information Theory. John Wiley (1991)

[7] Cowen, L., Ideker, T., Raphael, B.J., Sharan, R.: Network propagation: a universal amplifier of genetic associations. Nature Reviews Genetics 18(9), 551–562 (2017)

[8] Cyrus Maher, M., Bartha, I., Weaver, S., di Iulio, J., Ferri, E., Soriaga, L., Lempp, F.A., Hie, B.L., Bryson, B., Berger, B., Robertson, D.L., Snell, G., Corti, D., Virgin, H.W., Pond, S.L.K., Telenti, A.: Predicting the mutational drivers of future SARS-CoV-2 variants of concern. Science Translational Medicine 14(633), eabk3445 (2022)

[9] England, J.L.: Statistical physics of self-replication. J. Chem. Phys. 139, 121923 (2013)

[10] Fisher, R.A.: On the mathematical foundations of theoretical statistics. Philosophical Transactions of the Royal Society of London. Series A, Containing Papers of a Mathematical or Physical Character 222, 309–368 (1922)

[11] Fisher, R.A.: The Genetical Theory of Natural Selection. Clarendon Press, Oxford (1930). 10.5962/bhl.title.27468

[12] Gasteiger, J., Weißenberger, S., Günnemann, S.: Diffusion Improves Graph Learning. In: Wallach, H., Larochelle, H., Beygelzimer, A., d’Alché-Buc, F., Fox, E., Garnett, R. (eds.) Advances in Neural Information Processing Systems. vol. 32. Curran Associates, Inc. (2019)

[13] Hie, B., Zhong, E.D., Berger, B., Bryson, B.: Learning the language of viral evolution and escape. Science 371(6526), 284–288 (2021). 10.1126/science.abd7331

[14] Hie, B.L., Yang, K.K., Kim, P.S.: Evolutionary velocity with protein language models predicts evolutionary dynamics of diverse proteins. Cell Systems 13(4), 274–285.e6 (2022)

[15] Jordan, R., Kinderlehrer, D., Otto, F.: Free energy and the Fokker-Planck equation. Physica D 107, 265–271 (1997)

[16] Jordan, R., Kinderlehrer, D., Otto, F.: Variational formulation of the Fokker-Planck equation. SIAM J. Math Anal. 29(1), 1–17 (1998)

[17] Kimura, M.: Solution of a Process of Random Genetic Drift with a Continuous Model. Proceedings of the National Academy of Sciences of the United States of America 41(3), 144–150 (1955). 10.1073/pnas.41.3.144

[18] Kimura, M.: Diffusion Models in Population Genetics. Journal of Applied Probability 1(2), 177–232 (1964). 10.2307/3211856

[19] Koelle, K., Cobey, S., Grenfell, B., Pascual, M.: Epochal Evolution Shapes the Phylodynamics of Interpandemic Influenza A (H3N2) in Humans. Science 314(5807), 1898– 1903 (2006). 10.1126/science.1132745, https://www.science.org/doi/abs/10.1126/science.1132745

[20] Lawler, G.F.: Random Walk and the Heat Equation. American Mathematical Society, Providence, Rhode Island (2010)

[21] Lazarev, D.: Information measures for entropy and symmetry. 2211.14857 (2023)

[22] Sherman, M.A., Yaari, A.U., Priebe, O., Dietlein, F., Loh, P.R., Berger, B.: Genome-wide mapping of somatic mutation rates uncovers drivers of cancer. Nature Biotechnology 40(11), 1634–1643 (2022)

[23] Thadani, N.N., Gurev, S., Notin, P., Youssef, N., Rollins, N.J., Ritter, D., Sander, C., Gal, Y., Marks, D.S.: Learning from prepandemic data to forecast viral escape. Nature 622(7984), 818–825 (2023)

[24] Willett, J.D.S., Gravel, A., Dubuc, I., Gudimard, L., dos Santos Pereira Andrade, A.C., Lacasse, É., Fortin, P., Liu, J.L., Cervantes, J.A., Galvez, J.H., Djambazian, H.H.V., Zwaig, M., Roy, A.M., Lee, S., Chen, S.H., Ragoussis, J., Flamand, L.: Sars-cov-2 rapidly evolves lineage-specific phenotypic differences when passaged repeatedly in immune-naïve mice. Communications Biology 7(1), 191 (2024)

[25] Wright, S.: Evolution in mendelian populations. Genetics 16(2), 97–159 (mar 1931). 10.1093/genetics/16.2.97, https://academic.oup.com/genetics/article/ 16/2/97/6045152

[26] Yu, Y.W., Daniels, N.M., Danko, D.C., Berger, B.: Entropy-scaling search of massive biological data. Cell Systems 1(2), 130–140 (2023/06/11 2015). 10.1016/j.cels.2015.08.004, https://doi.org/10.1016/j.cels.2015.08.004

[27] Yubero, P., Manrubia, S., Aguirre, J.: The space of genotypes is a network of networks: implications for evolutionary and extinction dynamics. Sci. Rep. 7, 13813 (2017)

[28] Zadeh, L.A.: Fuzzy sets. Inf. Control 8, 338–353 (1965)

[29] Zadeh, L.A.: Probability measures of fuzzy events. J. Math. Anal. Appl. 23, 421–427 (1968)

[30] Zeng, T., Spence, J.P., Mostafavi, H., Pritchard, J.K.: Bayesian estimation of gene constraint from an evolutionary model with gene features. Nature Genetics 56(8), 1632–1643 (2024)

