## Appendix for "Constrained Diffusion as a Paradigm for Evolution"

### 1 Constrained diffusion equation derivation

We start with Eq. (8), writing it as,

$$\mathbf{f}(t) = \frac{1}{Z_t}\mathbf{k}(t) \circ (\mathbf{f}_0 e^{t\mathcal{L}}) = \frac{1}{Z_t}\mathbf{K}(t)(\mathbf{f}_0 e^{t\mathcal{L}}). \tag{1}$$

where

$$Z_t = \sum_{i=1}^{G} k_i(t)\,[\mathbf{f}_0 e^{t\mathcal{L}}]_i = \mathbf{1}_G^\top\,\mathbf{k}(t) \circ \mathbf{f}_0 e^{t\mathcal{L}}$$

is the normalization over the whole space $\mathcal{G}$ at time $t$, and $\mathbf{K}(t) := \mathrm{diag}(\mathbf{k}(t))$ is the $G \times G$ diagonal matrix with elements given by $\mathbf{k}(t)$. Writing $\mathbf{g}(\mathbf{t}) = \mathbf{f}_0 e^{t\mathcal{L}}$, we have, using a dot over the symbols to denote a time derivative, $\partial_t$,

$$\begin{aligned}
\dot{\mathbf{f}} &= \frac{1}{Z_t}\left(\dot{\mathbf{K}}\,\mathbf{g} + \mathbf{K}\,\dot{\mathbf{g}} - \frac{\dot{Z}_t}{Z_t^2}\mathbf{K}\,\mathbf{g}\right) \\
&= \frac{1}{Z_t}\left(\dot{\mathbf{K}}\,\mathbf{g} + \mathbf{K}\,\mathcal{L}\mathbf{g} - \frac{\dot{Z}_t}{Z_t}\mathbf{f}\right) \\
&= \frac{1}{Z_t}\left(\dot{\mathbf{k}} \circ \mathbf{g} + \mathbf{k} \circ \mathcal{L}\mathbf{g}\; - \mathbf{1}_G^\top(\dot{\mathbf{k}} \circ \mathbf{g} + \mathbf{k} \circ \mathcal{L}\mathbf{g})\;\mathbf{f}\;\right).
\end{aligned} \tag{2}$$

As in the main text, but in component form, we write $\mathcal{M}_i = (\mathcal{L}\mathbf{g})_i/\mathbf{g}_i$ and $\mathcal{S}_i = \dot{\mathbf{k}}_i/\mathbf{k}_i$. Writing (2) in component form, we have,

$$\begin{aligned}
\dot{\mathbf{f}}_i &= \frac{1}{Z_t}\left(\dot{\mathbf{k}}_i\mathbf{g}_i + \mathbf{k}_i(\mathcal{L}\mathbf{g})_i\; - \mathbf{f}_i\sum_{i=1}^{G}(\dot{\mathbf{k}}_i\mathbf{g}_i + \mathbf{k}_i(\mathcal{L}\mathbf{g})_i)\right) \\
&= \frac{1}{Z_t}\left(\mathbf{k}_i\mathcal{S}_i\mathbf{g}_i + \mathbf{k}_i\mathcal{M}_i\mathbf{g}_i\; - \mathbf{f}_i\sum_{i=1}^{G}(\mathbf{k}_i\mathcal{S}_i\mathbf{g}_i + \mathbf{k}_i\mathcal{M}_i\mathbf{g}_i)\right) \\
&= \mathbf{f}_i(\mathcal{S}_i + \mathcal{M}_i) - \mathbf{f}_i(\overline{\mathcal{S}}_i + \overline{\mathcal{M}}_i) \\
&= \mathbf{f}_i(\mathcal{S}_i - \overline{\mathcal{S}}_i) - \mathbf{f}_i(\mathcal{M}_i - \overline{\mathcal{M}}_i),
\end{aligned} \tag{3}$$

which gives, in vector form, Eq. (9).

### 2 Plots for Influenza Data

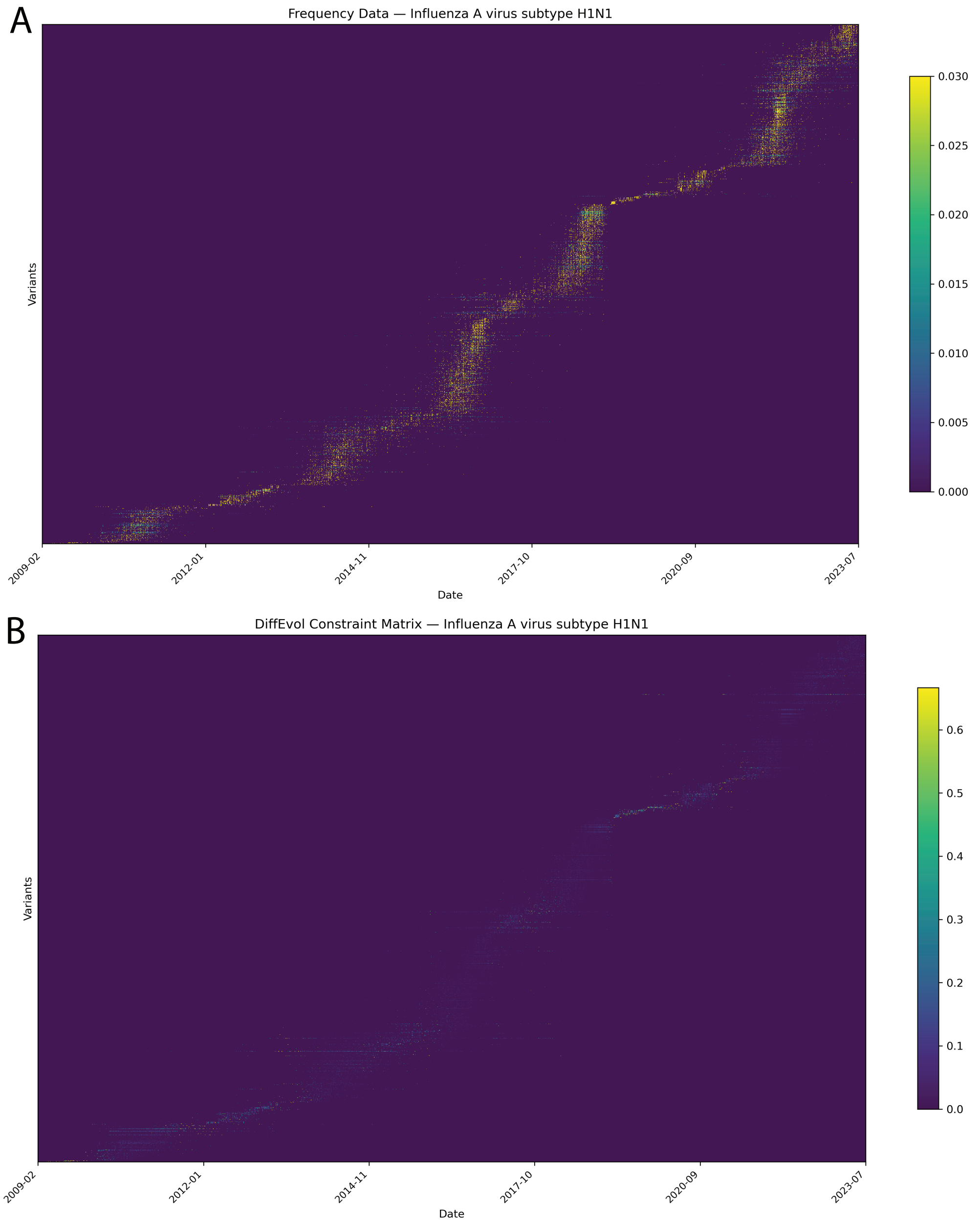


Figure 1: (A) heat map based on frequency data of the Influenza A virus (Subtype H1N1) nucleotide genotypes over twenty years. (C) Corresponding constraint matrix using `DiffEvol`. Variants are ordered by their mean emergence time.
